# Single nucleus RNA sequencing of juvenile dermatomyositis skeletal muscle identifies altered angiogenic signaling

**DOI:** 10.64898/2026.05.04.721749

**Authors:** Casey O. Swoboda, Carmy Forney, Cristofer Calvo, Lucinda P. Lawson, Hilal Cevik, Kairavee Thakkar, Caitlin Treuting, Stephen N. Waggoner, Cheryl Bayart, Meredith P Schuh, Alexander Zygmunt, Sheila T. Angeles-Han, Alexei Grom, Grant Schulert, Nathan Salomonis, Matthew T. Weirauch, Douglas P. Millay, Leah C. Kottyan, Shannon K. O’Connor

**Author notes:** **ORCID ID / email**: Casey O. Swoboda, BS, Carmy Forney, BS, Cristofer Calvo, PhD, Lucinda P. Lawson, PhD, Hilal Cevik, MD, PhD, Kairavee Thakkar, PhD, Caitlin Treuting, MS, Stephen N. Waggoner, PhD, Cheryl Bayart, MD, MPH, Meredith P. Schuh, MD, Alexander Zygmunt, MD, Sheila Angeles-Han, MD, Alexei Grom, MD, Grant Schulert, MD, PhD, Nathan Salomonis, PhD, Matthew T. Weirauch, PhD, Douglas Millay, PhD, Leah C. Kottyan, PhD, Shannon K. O’Connor, MD, PhD. **Senior corresponding author:** Shannon K. O’Connor, Address: Cincinnati Children’s Hospital Medical Center, 3333 Burnet Ave, Cincinnati, OH 45229, **Email:**.

## Abstract

Juvenile dermatomyositis (JDM) is a chronic multisystem vasculopathy and inflammatory myopathy characterized by proximal muscle weakness, distinct rash, and risk of complications such as calcinosis cutis, skin ulceration, and mortality. Molecular insight from diagnostic muscle biopsy histology is limited, and the mechanistic pathoetiology of JDM remains poorly defined. We used single nuclei transcriptomics to assess muscle samples from patients with newly diagnosed treatment-naïve JDM. As a control, we assessed muscle samples from patients with congenital (nemaline) myopathy (CM), a non-inflammatory disorder. A total of 25,794 high quality nuclei were analyzed and clustered into various muscle-resident or infiltrating cellular populations. JDM tissue was characterized by an enriched interferon (IFN) response signature across endothelial, stromal, and immune cell compartments. Endothelial and perivascular populations showed increased inflammatory and angiogenic programs. Intercellular communication inference analysis identified dysregulated vascular endothelial growth factor (VEGF)-related signaling involving endothelial, stromal, and myonuclear populations as a possible mechanism for myonuclear-driven modulation of the muscle microvasculature. Spatial RNA in situ hybridization supported increased expression of selected IFN responsive and angiogenesis signaling genes in JDM tissue. Collectively, these data provide a cell type-resolved view of treatment-naïve JDM muscle and highlight vascular and IFN pathways for follow-up in larger cohorts.

## Introduction

Juvenile dermatomyositis (JDM) is a rare multisystem vasculopathy and inflammatory myopathy characterized by proximal muscle weakness and distinct rash. Disease severity varies widely among patients with this condition. Severe symptoms include calcinosis cutis, dysphagia, and skin ulcerations^1^. In the United States, the annual incidence is 2 to 4 cases per 1 million children^2^, with a typical age of onset of 7 years^3–8^. JDM accounts for approximately 80% of idiopathic inflammatory myopathies in children. While the etiology of JDM remains unclear, JDM is considered an autoimmune reaction in genetically susceptible individuals, with environmental factors contributing to risk^3–8^.

Muscle pathology in JDM reflects microvascular injury and inflammation. Capillary loss, small-vessel changes, and immune infiltrates are common^9,10^. These findings support the concept that the muscle-endothelial interface is a key site of early injury. A consistent molecular hallmark of JDM is the increased expression of type I and type II interferon (IFN)-stimulated genes in blood and affected tissues^11–13^. IFN programs correlate with immune activation and may contribute to vascular injury^14^. However, the cellular sources of injury are only partially known. The relative contributions of infiltrating immune cells versus intrinsic changes in muscle and vascular cells also remain unclear^14–16^.

Recent transcriptomic studies have begun to define molecular programs in JDM. Single-cell peripheral blood mononuclear cell and whole blood analyses support IFN activation as a cardinal feature of this disease^11,12^. Spatial and single-cell analyses of muscle tissues from JDM patients corroborate the presence of IFN activation and immune cell activation, in addition to identifying mitochondrial dysfunction^17,18,19^. These studies identify candidate immune populations and tissue programs; however, cell-type-resolved mechanisms in treatment-naïve diagnostic muscle compared to non-inflammatory controls remain limited. Additionally, non-spatial single-cell analyses of skeletal muscle tissue exclude myofibers, the most abundant cell type in the tissue; their multi-nucleated structure requires single nuclei resolution.

Here, we used single nuclei RNA-seq profiling of treatment-naïve muscle biopsies to map muscle-resident and infiltrating populations in JDM. We assessed cell type-specific transcriptional programs with single-cell resolution, and identified the cellular dysregulation of angiogenesis in JDM, prioritizing pathways for follow-up. We included congenital (nemaline) myopathy (CM), allowing for a unique comparison between JDM tissue and tissue from a pediatric disease with a specific NEB genetic mutation known to cause physical shortening of the sarcomere apparatus, without an inflammatory component. Given that completely healthy children do not get muscle biopsies, this well-characterized comparator allows for focus on JDM specific features of abnormal muscle. Further, JDM samples and CM samples were collected and banked using the same protocols and processed in parallel, minimizing variation between samples. This study provides a cell type-resolved view of JDM diagnostic muscle, highlighting IFN and angiogenic programs, and reveals additional pathways beyond IFN that may be pathogenic in JDM.

## Results

### Clinicopathologic features of patients in this study

To anchor our molecular analyses, we first highlight characteristic clinical skin features of a treatment-naïve JDM patient, including the photosensitive dermatitis preferentially involving the upper eyelids (heliotrope), cheeks (malar), scalp, extensor joints (Gottron papules), and widespread vasculopathic changes on extensor surfaces, with complete sparing of flexural surfaces of the extremities^20^. More severe features, such as skin ulceration and contractures were also present in this patient (**Figure 1**). Additional clinical and lab results from the patients are summarized in **Table 1**.

**Figure 1.**
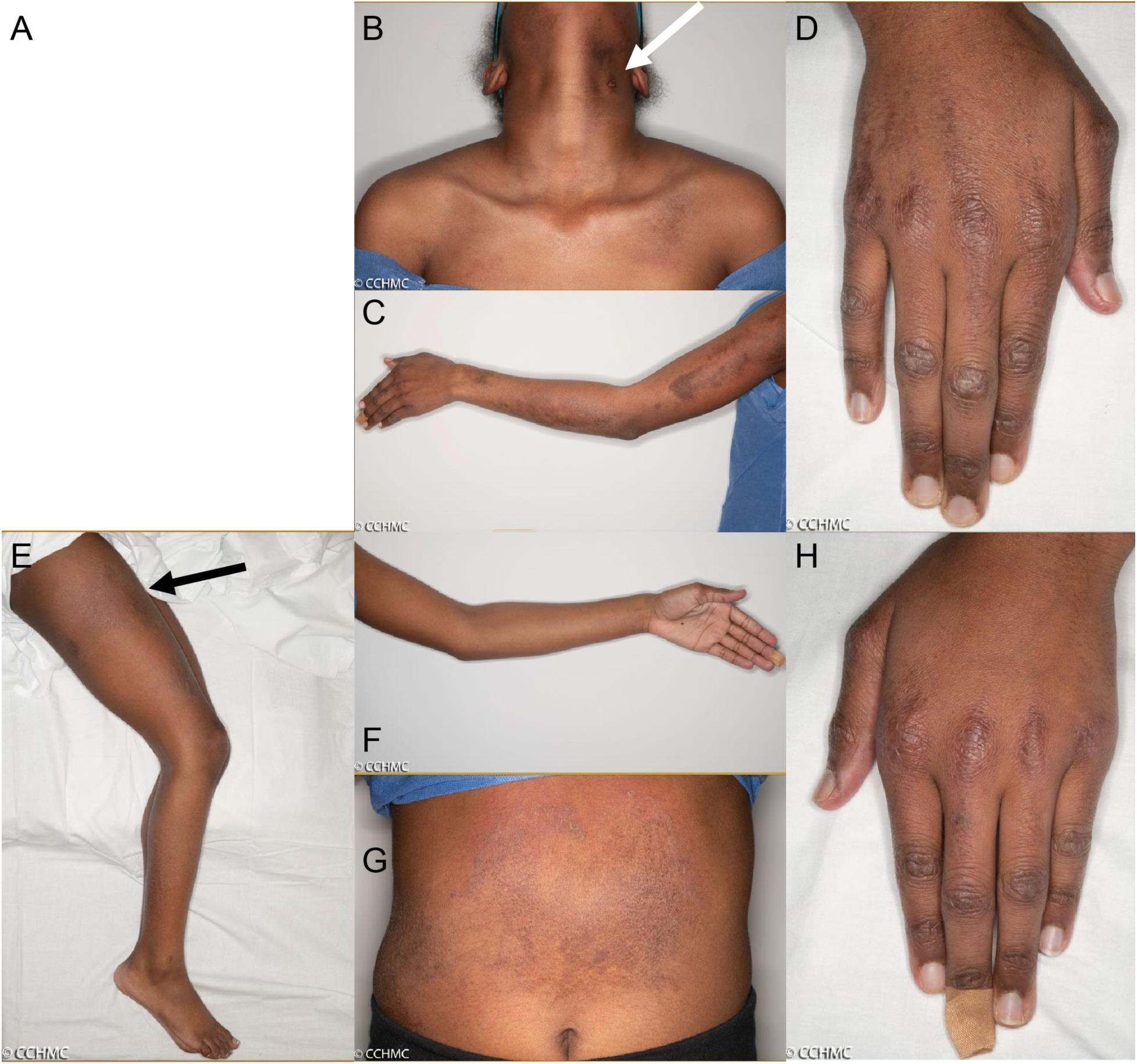
Skin and joint manifestations in juvenile dermatomyositis (JDM). Photosensitive dermatitis preferentially involving the A. upper eyelids (heliotrope rash), cheeks (malar rash), B. neck and chest (V sign), C. extensor surfaces, D-E. extensor joints (Gottron papules), F. lateral upper legs (Holster sign (arrow)), G. with complete sparing of flexural surfaces of the extremities. Less typical and more severe features identified on this patient include B. skin ulceration (arrow), C. contractures of wrists and elbows, H. abdominal vasculopathy. The patient depicted in this figure is included in the sequencing study presented in Figures 3-5.

**Table.** Patient characteristics. Patients with biopsies included in the single nuclei gene expression analysis are included in this table. CMAS, childhood myositis assessment scale; ID, identification; JDM, juvenile dermatomyositis; CM, congenital myopathy; NA, not applicable.

| *Characteristic* | *Value* |  |  |  |  |
| --- | --- | --- | --- | --- | --- |
| *Sample ID* | JDM1 | JDM2 | JDM3 | CM1 | CM2 |
| *Age at sample (y)* | 11.6 | 16.5 | 15.6 | 0.4 | 0.6 |
| *Sex* | F | F | F | F | F |
| *Myositis-specific antibody* | Mi-2 | TIF-1γ | Jo-1 | NA | NA |
| *JDM disease activity at time of biopsy:* |  |  |  |  |  |
| *CMAS (0-52)* | 1 | 38 | 18 | NA | NA |
| *CK, U/L* | 21,640 | 530 | 4103 | NA | NA |
| *LDH, U/L* | 2599 | 294 | 1029 | NA | NA |
| *ALT, U/L* | 546 | 45 | 66 | NA | NA |
| *AST, U/L* | 1380 | 90 | 120 | NA | NA |
| *ESR, mm/hr* | 52 | 9 | 28 | NA | NA |
| *CRP, mg/dL* | <0.40 | <0.29 | 1.4 | NA | NA |
| *Aldolase, U/L* | 258 | 13.1 | not collected | NA | NA |
| *Neopterin, nmol/L* | 22.3 | 30.6 | not collected | NA | NA |
| *Muscle Biopsy:* | Consistent with JDM | Consistent with JDM | Consistent with JDM | Consistent with nemaline myopathy | Consistent with nemaline myopathy |
| *Gene mutation* | NA | NA | NA | NEB gene mutation | NEB gene mutation |

Figure 2 illustrates the severity of the histopathologic features from the same patient. Muscle biopsy showed perifascicular myopathy with capillary bed abnormalities and interstitial fibrosis. Additional findings included severe intramuscular arteropathy, acute myopathic changes, focal myositis, the presence of muscle fibers with central nuclei, focal severe loss of capillary beds in areas of perifascicular endomysial fibrosis, and increased sarcolemmal MHC class I staining. Endothelial tubuloreticular inclusions were observed by electron microscopy, consistent with interferon exposure. These clinicopathologic features motivated our single nuclei profiling of diagnostic muscle to define cell type-specific programs in JDM.

**Figure 2.**
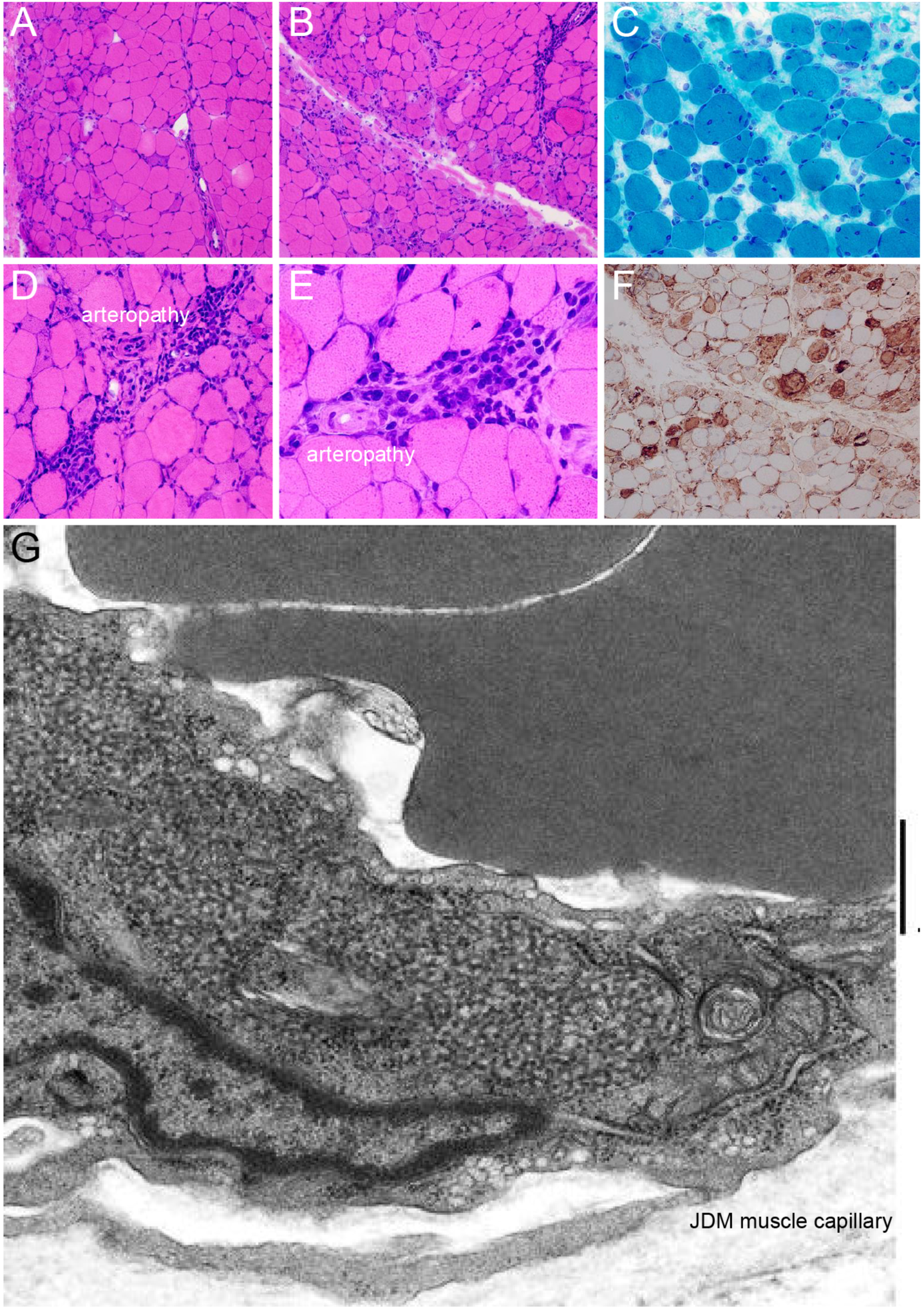
Representative histopathology and electron microscopy findings in muscle biopsy in severe juvenile dermatomyositis (JDM). A-B. Hematoxylin and eosin (H&E) stain. Severe peripheral fascicular myopathy. C. Trichrome stain. Interstitial fibrosis coincides with capillary loss. D-E. H&E stain. Arteropathy and focal myositis. F. MHC1 marks sarcolemma, necrotic fibers, and capillary bed changes at edge of fascicles. G. Electron microscopy. Tubuloreticular inclusions in endothelial cells reflect Interferon A production by endothelial cells. The patient sample depicted in this figure is included in the sequencing study presented in Figures 3-5. Images and interpretation by pathologist Dr. Kevin Bove.

### Single nuclei assessment of cell types

We generated single-nucleus transcriptomic profiles from treatment-naïve quadriceps biopsies from JDM (n=6) and congenital (nemaline) myopathy (CM, n=4) patients. These biospecimens were processed for genomic RNA expression analyses in parallel. After quality control (see **Supplemental Data Set** for QC information), we retained data from three JDM and two CM biopsies with 25,794 nuclei for analysis (Figure 3). We integrated data across donors and identified 19 transcriptionally distinct nuclear clusters (Figure 3A), which were variably represented in the biopsies from JDM (Figure 3B) and CM (Figure 3C) patient groups. We annotated clusters using canonical marker genes of various cell lineages (Figure 3D). Most nuclei mapped to expected muscle-resident compartments. These included multiple myonuclear states and myogenic progenitors. We also identified stromal and vascular populations, as well as immune cells. The overall composition was broadly similar across donors, regardless of diagnosis (Figure 3E).

**Figure 3.**
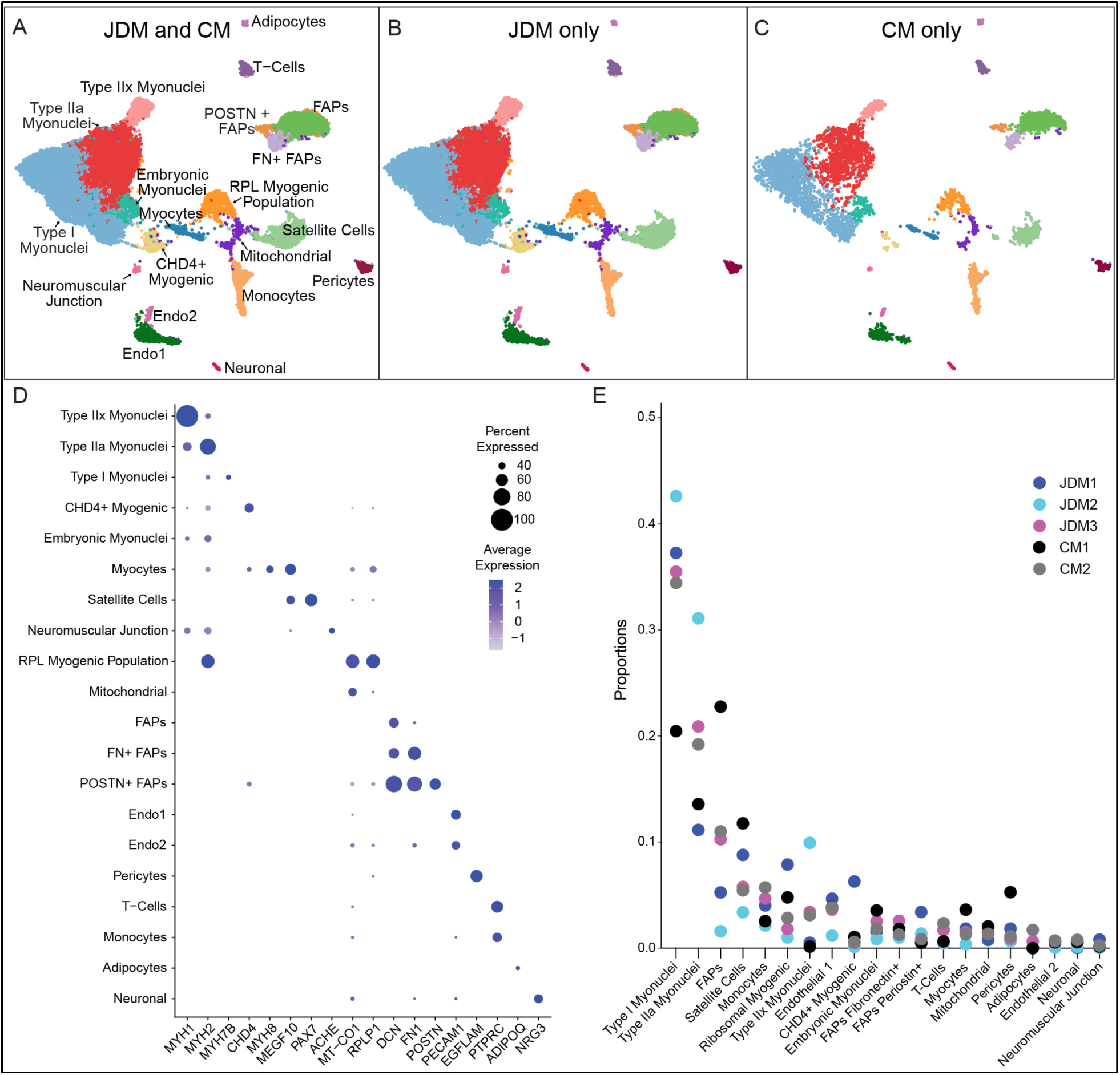
Single nuclei RNA of treatment naïve muscle biopsies from juvenile dermatomyositis (JDM, n=3) and comparator (congenital myopathy, CM, n=2). A. Integrated Uniform Manifold Approximation and Projection (UMAP) visualization representing 19 color-coded myogenic nuclei clusters identified by snRNA-seq in JDM and CM patient biopsies. Data were generated from quadricep muscles biopsies. B. Integrated UMAP of nuclei from JDM patient biopsies only. C. Integrated UMAP of nuclei from CM patient biopsies only. D. Dot plot showing expression levels of selected marker genes used to annotate major cell populations across clusters. E. Proportions of nuclei assigned to the clustered cell types for each JDM and CM patient.

We also observed a small cluster of myogenic nuclei with high *CHD4* expression (Figure 3B). This cluster was dominated by a single JDM donor with an anti-Mi-2 myositis specific antibody (MSA). Given the cohort size, we treat this as a donor-associated finding. Given that the CHD4 protein is the target of the Mi-2 autoantibody, it motivates follow-up in larger cohorts to test whether this state tracks with other patients with an anti-Mi-2 MSA, with serology, or with disease features.

### Pathways with JDM-specific differential expression across cell types

We next tested for JDM-associated transcriptional changes within each annotated cell population using pseudobulk differential expression analysis with donor as the unit of replication. So as not to overinterpret any one gene based on the smaller sample size, we used CellHarmony^21^ to perform pathway enrichment using Pathway Commons (see **Supplemental Data Set** for full results of CellHarmony pathway analysis). We visualized pathway signatures across cell types, allowing us to compare the direction and magnitude of differential expression of each gene set in JDM relative to CM (Figure 4).

**Figure 4.**
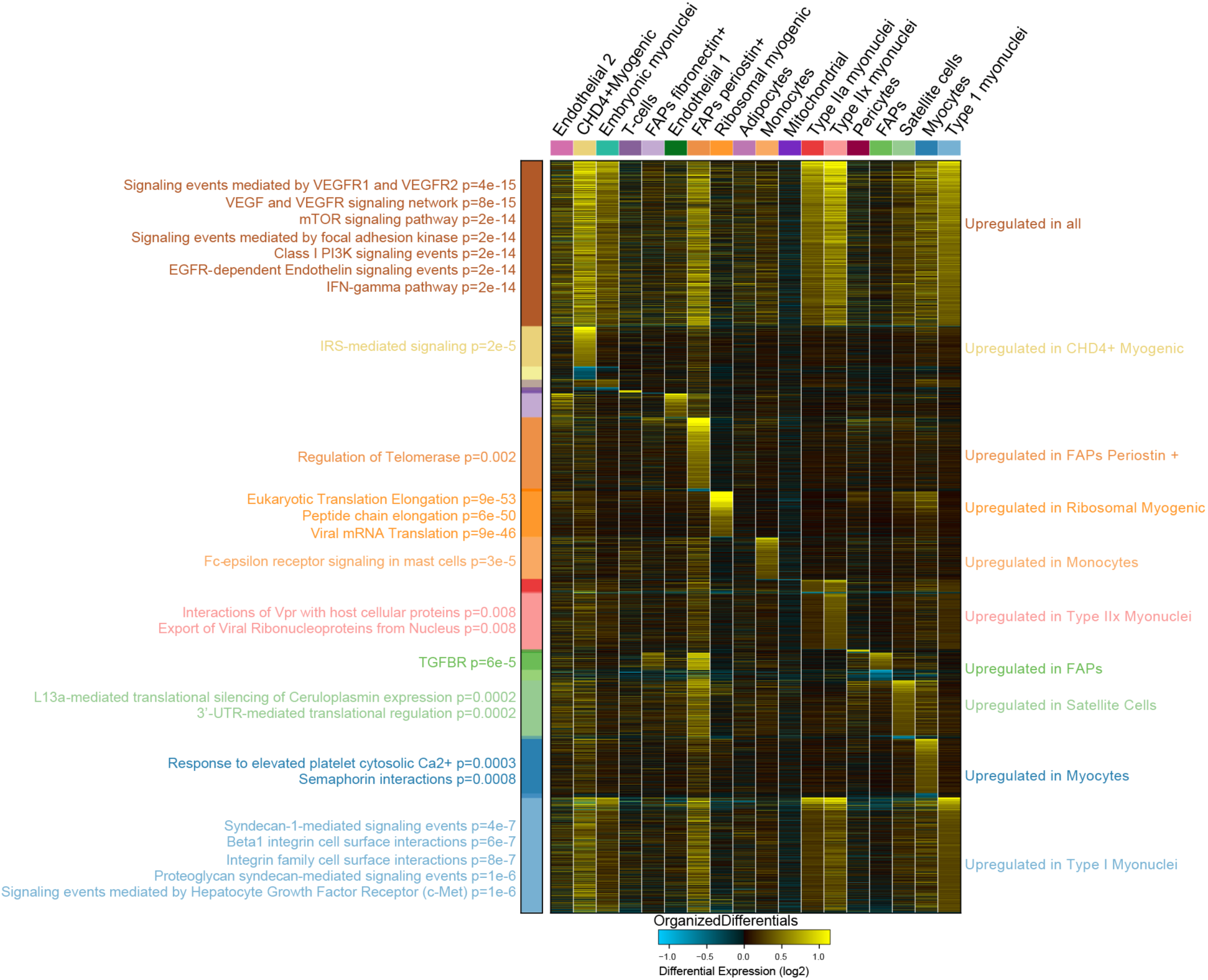
Pathways with JDM-specific differential expression across cell types. Differential expression analysis was performed within each annotated cell population using donor-level pseudobulk profiles comparing juvenile dermatomyositis (JDM; query, n=3) to congenital (nemaline) myopathy (CM; reference, n=2). Empirical Bayes moderated t-tests were used with Benjamini-Hochberg false discovery rate (FDR) correction. Differentially expressed genes were defined by fold change >1.2 and FDR-adjusted p<0.05. Differential genes were grouped based on whether they were significant across multiple cell populations or restricted to a single population. Pathway enrichment was performed using the Pathway Commons database. The CellHarmony heatmap displays log2 differential expression in JDM relative to CM across cell populations (see color scale). Fisher’s exact test P-values shown next to pathway terms are from the Pathway Commons enrichment step and indicate statistically significant gene sets. See Supplemental Dataset 1 for full results.

Several pathway signatures were increased across many compartments in JDM, including the VEGF and VEGF-receptor signaling (*VEGFR1*/*VEGFR2*) pathways (Figure 4). Although VEGF upregulation is a known feature of skeletal muscle in adult myositis^22,23^, our data reveal an unexpected breadth of this VEGF signature across many distinct cell types within the biopsy tissue in JDM. We also observed enrichment of signaling pathways annotated to mTOR, focal adhesion kinase, class I PI3K, and EGFR-dependent endothelin signaling, some of which have been implicated before in autoimmune conditions^24–26^. Additionally, we confirm the presence of prominent interferon-associated programs across multiple cell populations, including enrichment of an IFN-γ-annotated pathway gene set (Figure 4).

In addition to shared pathway signatures, we also observed cell type-restricted pathway signatures. IRS-mediated signaling was most evident in the CHD4+ myogenic cluster (specific to the JDM donor with Mi-2 autoantibodies). Regulation of telomerase was enriched in periostin expressing fibro-adipogenic progenitors (FAPs). A ribosomal myogenic cluster showed strong enrichment of translation-related gene sets. Monocytes showed enrichment of an Fcε receptor-annotated pathway gene set. Type IIx myonuclei showed enrichment of pathways associated with viral nuclear export and ribonucleoprotein trafficking. Satellite cells showed enrichment of 3′-UTR-mediated translational regulation and L13a-mediated translational silencing. Myocytes showed enrichment of semaphorin interactions and a platelet cytosolic Ca2+-response-annotated term. The type I myonuclei showed enrichment of extracellular matrix and adhesion-associated signaling, including syndecan/integrin and c-Met-annotated pathways (Figure 4). Each of these cell type-restricted pathway signatures warrant further in-depth investigation.

Together, these analyses highlight broad vascular and IFN-associated programs spanning multiple cell types in diagnostic JDM muscle. They also nominate distinct myogenic and stromal pathways for targeted analysis and mechanistic study in larger cohorts.

### Cell-cell communication inference of ligands and receptors highlights JDM-specific dysregulation of VEGF signaling

To nominate candidate signaling axes that could link vascular and stromal changes in JDM muscle, we performed cell-cell communication inference using CellChat.^27^ In CM, inferred VEGF communication probabilities were highest as autocrine signaling within endothelial cells, as well as signaling from FAPs to endothelial cells, as expected^28^ (Figure 5A). In contrast, JDM showed a distinct pattern of inferred VEGF signaling, with increased communication probabilities across multiple muscle-resident and stromal populations, but comparatively less autocrine signaling within endothelial cells (Figure 5B). Dysregulation of the VEGF signature identified in pathway and cell communication analyses (**Figures 4 and 5)** are consistent with prior immunohistochemical histologic findings of increased VEGF protein expression^22,23^. Network visualization of aggregated VEGF signaling highlighted myogenic induction of VEGF programs in our JDM group, absent in our comparator group, CM (Figure 5C). In JDM, endothelial and stromal compartments contributed prominently to inferred VEGF signaling, with predicted interactions involving endothelial subpopulations (Endo1/Endo2), FAPs, and myonuclear populations (Figure 5D). These results, based on transcriptomic inference, nominate VEGF-directed communication as a pathway for targeted analysis and mechanistic study; namely, these results invite an explanation for the dichotomy of profound angiogenic signaling without evidence of new vascular formation.

**Figure 5.**
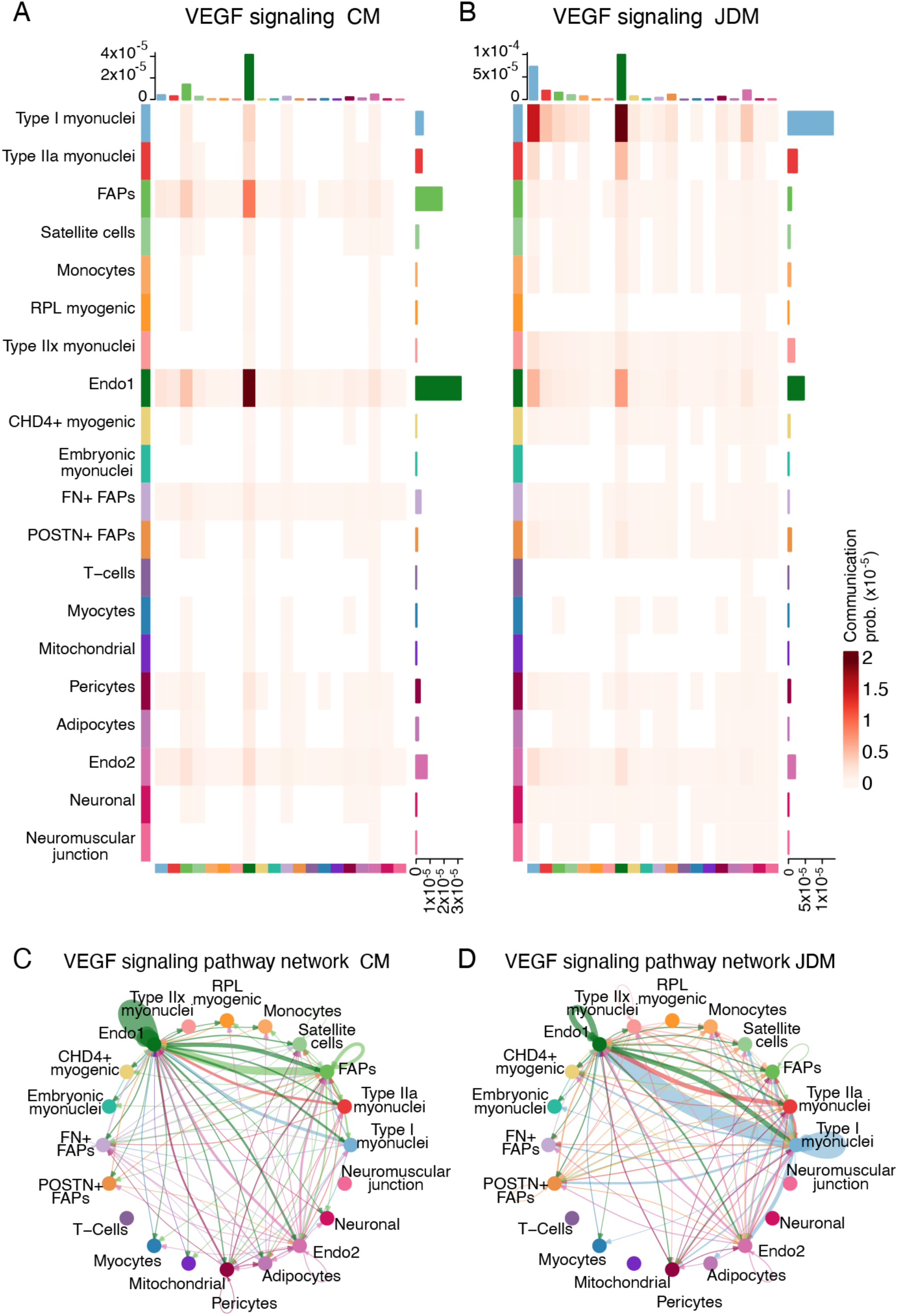
Cell-cell communication of ligands and receptors in the VEGF pathway. A-B. The inferred communication of VEGF signaling molecules from cells identified in this study (see **Figure 3**). Shading of boxes within each subpanel is a function of the communication probability (darker shading indicates higher probability, see scale legend). Rows indicate sender (source) cell populations and columns indicate receiver (target) populations. Color intensity reflects the relative communication probability. Side bars show total outgoing signaling strength by sender population. Top bars show total incoming signaling strength by receiver population (as computed by CellChat). A. Congenital myopathy (CM, n=2). B. Juvenile dermatomyositis (JDM, n=3). C-D. Aggregated VEGF signaling networks inferred by CellChat for CM (C) and JDM (D), generated independently. Nodes represent cell populations. Edge width reflects inferred interaction strength. Edge color denotes the sender population.

### Spatial RNAscope of specific RNA expression

To provide orthogonal, spatial support for key transcriptional inferences from the single nuclei dataset, we performed RNAscope on muscle sections and examined transcripts prioritized from the differential expression and communication analyses. We focused on *VEGFA*, *CXCL9*, and *VWF* (von Willebrand factor), which index vascular signaling, IFN-associated chemokine programs, and endothelial disruption, respectively.

Across evaluated sections, JDM muscle showed a qualitatively increased *VEGFA* signal compared to controls (Figure 6). Signal was most apparent in perivascular and stromal regions, consistent with increased vascular signaling programs identified in the transcriptomic analyses. JDM tissue also showed qualitatively increased *CXCL9* signal, consistent with an IFN-associated inflammatory milieu in diagnostic muscle (Figure 6). Finally, *VWF* signal was sparsely identified in all tissue sections (Figure 6). We note that these RNA-FISH findings support spatial localization of the selected transcripts rather than providing quantitative validation.

**Figure 6.**
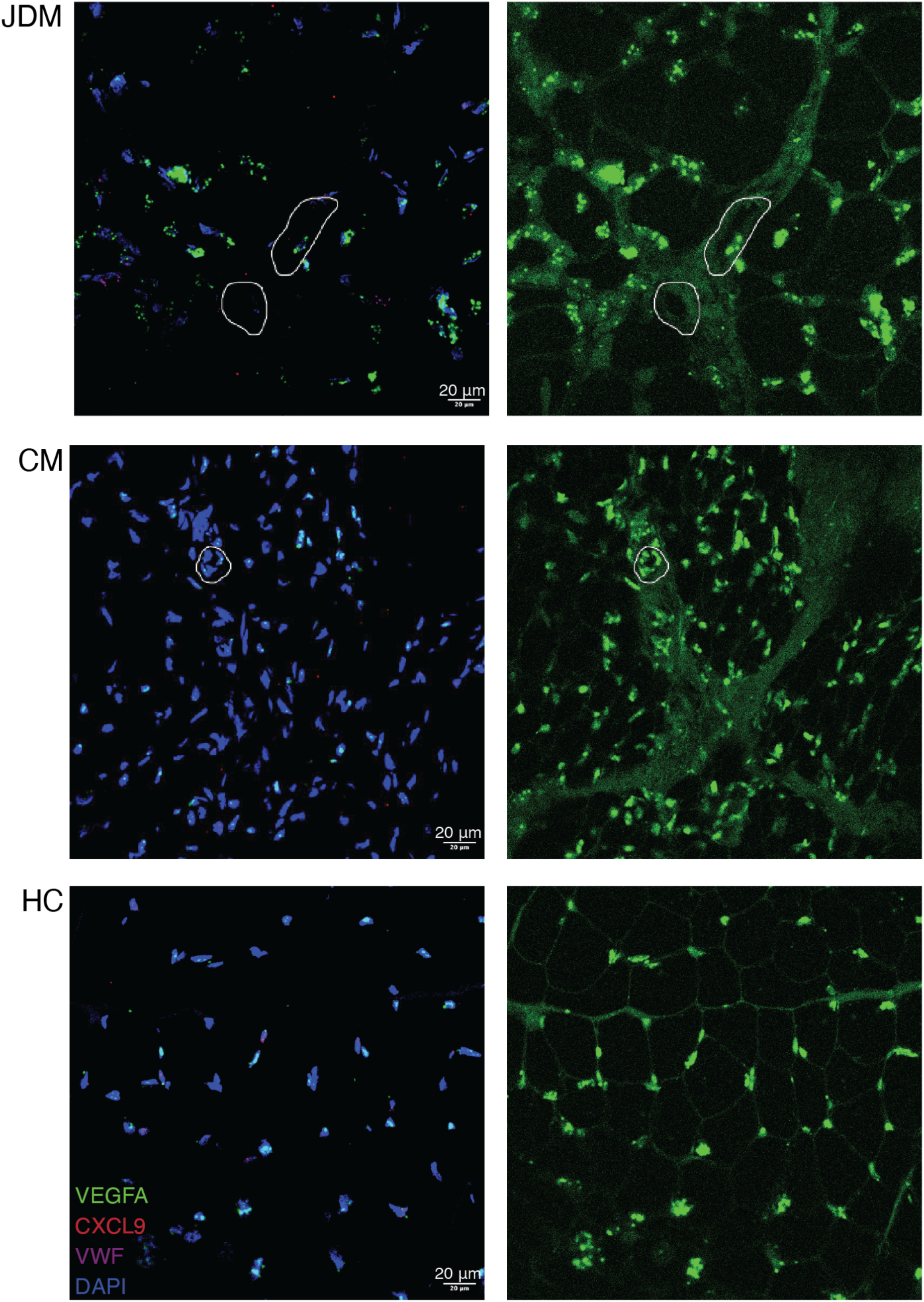
Spatial RNAscope of specific RNA expression. Representative confocal microscopy images of spatial RNA fluorescence in situ hybridization (RNA-FISH) images for specified genes of sections from A. Juvenile dermatomyositis (JDM), B. Congenital nemaline myopathy (CM) C. Healthy control (HC). Signal for *VEGFA* (green), *CXCL9* (red), and *VWF* (magenta) overlaid in the left column. Nuclei counterstained with DAPI (blue). Images illustrate increased qualitative VEGFA and CXCL9 signal in JDM muscle relative to CM and HC, with prominent signal in perivascular regions. Scale bars are indicated on each figure panel. Images in the right column demonstrate GFP autofluorescence of the sections, in addition to *VEGFA* expression, to identify the location of fascial planes and blood vessels. Dashed white lines are provided to indicate blood vessels in the representative image from JDM and CM. Blood vessels were not identified in captured sections from HC. Findings are qualitative and intended to support spatial localization of selected transcripts. Images captured at 20x magnification.

## Discussion

In this pilot study, we profiled treatment-naïve diagnostic muscle biopsies from children with juvenile dermatomyositis (JDM), with a non-inflammatory congenital (nemaline) myopathy comparator processed in parallel. We used single nuclei transcriptomics to map muscle-resident and infiltrating populations. We then performed cell type-resolved differential expression analysis. We prioritized pathway-level signals to limit over-interpretation of individual genes in a small cohort. Two important themes emerged. First, IFN-associated programs were detectable across multiple compartments. In addition to confirming an IFN signature in JDM, our results provide an original demonstration of more widespread IFN responses across multiple cell types in skeletal muscle than was previously appreciated. Second, vascular and endothelial signaling pathway signatures were prominent and converged on VEGF-related pathways, similarly with more widespread VEGF-ligand RNA expression across cell types than previously identified.

Interferon-stimulated gene programs are a reproducible hallmark of JDM in blood and tissues, and our data are consistent with that literature^12^. Our results extend these observations by localizing IFN-associated signatures across vascular, stromal, and immune compartments in diagnostic muscle. We also observed *CXCL9* signal by RNAscope, which is consistent with an IFN-primed inflammatory milieu within the tissue^12^. These findings complement recent single-cell profiling efforts in JDM that emphasize coordinated immune dysregulation and type I IFN activation across immune populations^12^.

A second key finding was the prominence of angiogenic signaling from JDM muscle. JDM is defined clinically and pathologically by vasculopathy, and prior work has linked microvascular injury to impaired angiogenic responses and IFN-induced angiostatic chemokines in JDM muscle^14^. In adult inflammatory myopathies, *VEGF* has been reported to be elevated in muscle, and the *VEGF* isoform balance may shift toward anti-angiogenic forms^22,23^. In our dataset, pathway enrichment analysis highlighted altered VEGF signaling and related endothelial signaling pathway signatures across multiple cell populations. Cell-cell communication inference further nominated aberrant VEGF-directed signaling as a candidate axis linked to the profound vasculopathy of JDM. These analyses do not demonstrate functional signaling. They provide a hypothesis-generating framework that suggests that the VEGF network may represent either a compensatory response to microvascular injury, a maladaptive pathway, or both. RNAscope provided qualitative spatial support for *VEGFA* signals in diseased tissue. These observations motivate targeted follow-up in larger cohorts with quantitative spatial assays and orthogonal protein-level measurements in future studies.

Our findings should be interpreted in the context of recent single-cell and spatial transcriptomic studies of JDM muscle. A prior single-cell RNA-seq study profiled peripheral blood and muscle and reported inflammatory features in JDM muscle pre and post treatment^19^. Spatial approaches have also highlighted heterogeneity within muscle and persistence of abnormal programs over time^18^. A recent treatment-naïve spatial transcriptomics study identified robust interferon signatures and also emphasized mitochondrial abnormalities^17^. We did not evaluate mitochondrial pathology in a systematic way here. Our work adds complementary resolution by combining donor-aware cell type-specific differential expression with ligand-receptor inference to prioritize candidate cross-compartment signaling axes.

This study has important limitations. The primary limitation is sample size. Only three JDM biopsies and two comparator biopsies yielded sufficient quality for the sequencing analysis. This limits power to detect subtle composition shifts while increasing sensitivity to donor-specific effects. We therefore treated donor as the independent unit of replication (i.e.. pseudobulked) for differential expression and emphasized pathway-level patterns. Comparator choice is another limitation. Given that healthy children do not receive muscle biopsies, thoughtful consideration of comparator group led us to Nemaline myopathy, with a specific and known gene mutation affecting the length of the sarcomere, the unit of muscle contraction. Nemaline myopathy is non-inflammatory, but it is not a normal muscle baseline. Diagnostic muscle biopsies in these patients occurred in infancy, rather than school-age patients, as was the case for the JDM patients. Some transcriptional differences may reflect distinct muscle remodeling states and the age of patients at diagnostic biopsy date rather than JDM-specific biology.

Additionally, by chance, the five biopsies (JDM and CM) with nuclei that passed quality assessment were all female. JDM patients are predominantly female with a ratio of 2:1^3–8^, but an all-female cohort may impact translatability to the general population with this diagnosis. Cell-cell communication analyses are useful but are based on inferences that depend on expression levels, curated ligand-receptor databases, and model assumptions. The RNAscope analyses were qualitative and not powered for statistical comparison. Finally, although nuclei were generated using a multiome workflow, the present manuscript focuses exclusively on the transcriptomic layer as ATAC data did not meet QC thresholds. Integration of chromatin accessibility to link cell type-specific regulatory elements to gene programs is a key next step.

This study provides a cell type-resolved view of treatment-naïve JDM diagnostic muscle. It supports interferon-associated programs across multiple compartments and highlights vascular pathway signatures that converge on VEGF-directed communication. These results generate testable hypotheses for larger studies that integrate spatial quantification, proteomics, and regulatory genomics. Future work should test whether the programs identified here stratify by myositis-specific autoantibodies or clinical phenotype. This includes analysis of donor-associated signals, such as the CHD4-high myonuclear state observed in one anti-Mi-2-positive case. Larger cohorts will also enable formal integration of transcriptional programs with histopathologic scoring of severity and chronicity along with quantitative measures of microvascular density and injury. These studies will be necessary to determine whether VEGF-related pathways represent treatable biology, compensatory repair, or a marker of disease stage.

## Methods

### Sex as a biological variable

Both male and female participants were included, and by chance, the five biopsies with nuclei that passed quality assessment were all female. Sex is reported for each participant in Table 1.

### Patient recruitment

This study was approved by the Cincinnati Children’s Hospital Medical Center (CCHMC) Institutional Review Board (protocol 2017-0430). Written informed assent and/or consent was obtained from patients and parents before participation.

We enrolled children undergoing diagnostic muscle biopsy who were treatment-naïve at CCHMC. JDM was defined using the Bohan and Peter criteria^1,29^, with histopathological confirmation from muscle biopsy. Congenital (nemaline) myopathy (CM) was used as a non-inflammatory myopathy comparator. CM diagnosis was established by histopathology and supported by identification of pathogenic variants in NEB (nebulin, chromosome 2)

In total, we enrolled 6 JDM participants and 4 CM participants. Single nuclei sequencing analyses were performed on the subset of biopsies that yielded sufficient nuclei quality and quantity. This primary sequencing cohort included 3 JDM biopsies and 2 CM biopsies.

Additional JDM (n=3) and CM (n=2) biopsies were processed but did not meet quality thresholds for inclusion in the primary sequencing analysis.

RNAscope analyses were performed on available biopsy sections, including the 6 JDM patients, the 4 CM patients, and 4 ‘healthy controls’ (HC) patients. These 4 HC patients were evaluated for muscle weakness and underwent muscle biopsy but did not receive a specific diagnostic classification and did not have evidence of inflammation. These biopsies were used for exploratory RNAscope imaging only.

### Muscle biopsies

Muscle biopsies were obtained from treatment-naïve patients with JDM, CM, and HC. Biopsies were wrapped in foil and flash-frozen in liquid nitrogen within 30 minutes of collection. Frozen tissue was transferred to cryovials and stored at −80°C in the CCHMC Pathology Core until processing.

Mouse muscle was collected as a technical control for workflow optimization. Tibialis anterior and gastrocnemius muscles were harvested from C57BL/6 mice euthanized under IACUC-approved protocols for unrelated studies. Tissue was flash-frozen in liquid nitrogen immediately after harvest and stored at −80°C. Mouse samples were used for nuclei isolation and library preparation optimization and were not used for JDM–CM comparisons.

### Isolation of nuclei

Single nuclei were isolated from frozen muscle using the 10x Genomics protocol “Nuclei Isolation from Complex Tissues for Single Cell Multiome ATAC + Gene Expression Sequencing.”

All steps were performed on ice or at 4°C. Frozen tissue was mechanically dissociated and lysed to release nuclei. Nuclei were washed and resuspended in nuclei buffer for downstream multiome processing.

Protocol optimization was performed using murine muscle. During optimization, we found that nuclei quality and yield were sufficient without fluorescence-activated nuclei sorting. Therefore, sorting was not performed for patient samples. Each patient biopsy was processed individually. For each multiome run, a mouse muscle nuclei preparation was processed in parallel as a technical control. Immediately prior to Chromium loading, we added 10% murine nuclei to the human nuclei preparation as a species-mixing run control. Murine-dominant nuclei were excluded during downstream processing and were not used for analyses presented in this manuscript.

### Single nuclear RNA and ATAC-sequencing (snRNA-seq and snATAC-seq)

Single nuclei RNA sequencing (snRNA-seq) and single nuclei ATAC sequencing (snATAC-seq) libraries were generated using the Chromium Next GEM Single Cell Multiome ATAC + Gene Expression workflow (10x Genomics). Following nuclei isolation, nuclei were counted using a hemocytometer. Nuclei concentration was adjusted to fall within the range recommended by the manufacturer for Chromium loading. Up to 15,000 nuclei were then loaded per reaction onto the Chromium chip and processed according to the manufacturer’s protocol. Library preparation was performed by the CCHMC Single Cell Genomics Facility.

### Data processing

Initial read alignment and quantification of multiome fastq files were generated using CellrangerARC/v2.02, aligned to the GRCh38-2020-A genome build provided by 10X Genomics. For each dataset, generated BAM files were separated into RNA and ATAC reads using the bam2fastq function provided by cellranger. RNA filtered fastq files were aligned to a combined genome reference of GRCh38 and GRCm39 provided by 10X genomics, and quantified using Cellranger/v7.0.1 count, setting the –-chemistry parameter to “ARC-v1”. Cellbender/v0.3.2 was used to remove ambient mRNA from the raw count matrices output. Cellbender corrected matrices were then imported into seurat objects (Seurat/5.1.0). Cells were identified as “Mouse” if greater than 90% of their counts aligned to reads in the mouse genome. These were then removed from all downstream analyses. The count matrices were then filtered to remove all rows containing genes from the mouse genome. Nuclei with greater than the 98^th^ percentile of unique features, less than 200 features, or greater than 5% of reads mapping to the mitochondrial genome were also removed from downstream analysis. Seurat objects with the remaining nuclei then underwent doublet identification using Solo/v0.2. Seurat objects were then merged, with normalization, variable feature selection, and scaling performed using SCTransform(). Principal component analysis was performed using the RunPCA() function. ATAC outputs were generated by Cell Ranger ARC but were not analyzed further for this study due to data quality in some samples.

### snRNA-seq data integration

The merged Seurat object was integrated using the IntegrateLayers() function, specifying the “method” argument as HarmonyIntegration and the normalization.method argument as “SCT”. Further dimensionality reduction was performed using FindNeighbors(), FindClusters(), and RunUMAP(), using thirty principal components and a resolution of 0.5. Doublets were then filtered from the object, and an additional round of dimensionality reduction was performed, using a resolution value of 1. Clusters were then merged and annotated based on the expression of known cell type markers.

### Pathway analyses

Differential expression (DE) analysis was performed within each annotated cell population using CellHarmony. For each donor and cell type, we generated pseudobulk expression profiles and compared the query group (JDM) to the reference group (CM). DE was tested using an empirical Bayes moderated t-test with Benjamini-Hochberg false discovery rate (FDR) correction. Genes were considered differentially expressed at fold change >1.2 and FDR-adjusted p<0.05. Differentially expressed genes were grouped based on whether they were significant in a single cell population or across multiple populations. Gene set enrichment analysis was then performed for each gene group using the Pathway Commons database. Pathway-level signatures and organized heatmaps were generated using CellHarmony,^21^ version 2, run August 2025. In parallel, CellHarmony also reports per-gene t-tests treating cells as replicates; donor-aware pseudobulk results were used for inference.

### Cell-cell communication analyses

Cell-cell communication analysis was performed using CellChat (v2.1.2). Log-normalized gene expression levels from the integrated Seurat object were split by diagnosis (JDM or CM), while retaining the integrated cell-type labels. For each condition, CellChat identified overexpressed genes and predicted ligand-receptor interactions using the built-in human CellChat database. Communication probabilities were computed using the tri-mean method, with adjustment for population size, and were smoothed using the human protein-protein interaction (PPI) network. Predicted interactions were filtered to retain signaling between cell groups containing at least 3 cells. To compare conditions, CellChat objects were harmonized across shared cell identities and merged. Differences in VEGF signaling were visualized using netVisual_heatmap(), and aggregated VEGF signaling networks were visualized using netVisual_aggregate().

### Spatial RNA-FISH of RNA expression

Spatial RNA-FISH was performed using ACD-Bio RNAscope Multiplex Fluorescent Reagent Kit v2, following manufacturer’s instructions. Probes were designed for *VEGFA*, *CXCL9*, and V*WF*, with DAPI counterstain, as well as negative and positive assay controls provided by the manufacturer for each slide. 3 images per muscle biopsy section were captured at 20x magnification from 6 patients with JDM, 4 patients with CM, and 4 HC patients (see Patient Recruitment section for clinical descriptions). Imaging was performed on a Nikon a1plus Confocal microscope. Image compilation was performed in Fiji.^30^

### Statistics

Clinical characteristics are presented descriptively. For single-nucleus RNA-seq analyses, donor was treated as the independent unit of replication for differential expression analyses within each annotated cell population. Pseudobulk expression profiles were generated for each donor and cell type and compared between JDM and CM using CellHarmony with empirical Bayes moderated t-tests and Benjamini-Hochberg FDR correction. Genes were considered differentially expressed at fold change greater than 1.2 and FDR-adjusted P less than 0.05.

Pathway enrichment analyses were performed in CellHarmony using the Pathway Commons database, and Fisher exact test P values are shown for enriched gene sets. Cell-cell communication analyses performed with CellChat were used for inference of putative ligand-receptor signaling networks and were not intended as formal statistical proof of functional signaling. RNAscope findings were evaluated qualitatively and were not powered for formal statistical comparison. No additional statistical methods were used.

### Study approval

All human studies were approved by the Institutional Review Board of Cincinnati Children’s Hospital Medical Center (protocol 2017-0430). Written informed consent and/or assent was obtained from participants and their parents or legal guardians prior to participation. The manuscript includes a clinical photograph of a patient with juvenile dermatomyositis because the image documents disease manifestations that are important to the clinical message, including findings in dark skin that are underrepresented in the teaching and scholarly literature. Written informed consent for publication of the photograph was obtained and the record of consent has been retained in the electronic medical record and is available upon request. Animal tissue used for technical workflow optimization was collected under Institutional Animal Care and Use Committee approval at Cincinnati Children’s Hospital Medical Center.

### Data availability

Raw sequencing data has been deposited in GEO under accession number GSE329054 and are publicly available. Source data are provided.

### Code availability

Example of what goes here: Data analysis is described in the Methods. Scripts are available on GitHub at https://github.com/cswoboda/JDM-snRNAseq/issues

## Author contributions

CS contributed to conceptualization, methodology, data curation, formal analysis, visualization, and writing (original draft and editing). CF contributed to methodology, investigation, and writing (editing). CC contributed to investigation. LPL contributed to visualization and writing (editing). HC contributed to sample acquisition. KT contributed to methodology. CT contributed to visualization. SNW contributed to writing (editing) and supervision. CB contributed to clinical expertise and expert patient phenotyping. MS contributed to methodology and writing (editing). AZ contributed to writing (editing), clinical expertise, and expert patient phenotyping. SAH contributed to writing (editing), supervision, and clinical expertise. AG contributed to writing (editing), supervision, clinical expertise, and funding acquisition. GS contributed to methodology, writing (editing), supervision, and clinical expertise. NS contributed to methodology, formal analysis, and supervision. MTW contributed to conceptualization, methodology, formal analysis, writing (editing), supervision, and funding acquisition. DPM contributed to conceptualization, methodology, formal analysis, writing (editing), supervision, and funding acquisition. LCK contributed to conceptualization, methodology, sample acquisition, formal analysis, visualization, writing (original draft and editing), supervision, and funding acquisition. SKO contributed to conceptualization, methodology, sample acquisition, investigation, data curation, formal analysis, visualization, writing (original draft and editing), supervision, clinical expertise, expert patient phenotyping, and funding acquisition.

## Funding Support

SKO was supported by NIH T32 AR069512, Cure JM Fellowship Award, and the Cincinnati Children’s Research Foundation (CCRF). MTW and LCK were supported by NIH P30 AR070549 and CCRF ARC Award #53632. This work is the result of NIH funding, in whole or in part, and is subject to the NIH Public Access Policy. Through acceptance of this federal funding, the NIH has been given a right to make the work publicly available in PubMed Central.

## Acknowledgements

We dedicate this work to Kevin E. Bove, MD, who served the Cincinnati Children’s Hospital community as a pathologist and scientist for 56 years. Fascinated by juvenile myositis his whole career, he spearheaded the now internationally validated scoring system predicting chronicity and severity from muscle biopsies in children with JM. As much as he illuminated about this disease, there were questions he could not answer about the nature of the interplay between the vascular pathology and the muscle dysfunction in these patients with methods available at the time. He carefully and consistently preserved these samples and was the pathologist of record for every muscle biopsy for every child at CCHMC for his entire career. Dr. Bove inspired and helped design this study before he passed away. We hope we have taken the first step to answer his questions.

This research was made possible, in part, using the Cincinnati Children’s Single Cell Genomics Facility [RRID:SCR_022653], the Cincinnati Children’s Integrated Pathology Research Facility [RRID: SCR_022637], and the Cincinnati Children’s Imaging Research Center [RRID: SCR_022631].

## Notes

### Competing Interest Statement

The authors have declared no competing interest.

